# Single molecule occupancy patterns of transcription factors reveal determinants of cooperative binding *in vivo*

**DOI:** 10.1101/2020.06.29.167155

**Authors:** Can Sönmezer, Rozemarijn Kleinendorst, Dilek Imanci, Laura Villacorta, Dirk Schübeler, Vladimir Benes, Arnaud R Krebs

## Abstract

Gene activation requires the cooperative activity of multiple transcription factors at cis-regulatory elements. Yet, most transcription factors have short residence time, questioning the requirement of their physical co-occupancy on DNA to achieve cooperativity. Here, we advance Single Molecule Footprinting to detect individual molecular interactions of transcription factors and nucleosomes with DNA at mouse cis-regulatory elements. We apply this strategy to quantify the simultaneous binding of multiple transcription factors on single DNA molecules. Analysis of the binary occupancy patterns at thousands of motif combinations reveals that for most types of transcription factors high DNA co-occupancy can occur in absence of direct physical interaction, at sites of competition with nucleosomes. Perturbation of pairwise interactions demonstrates the function of molecular co-occupancy for binding cooperativity. These findings elucidate the binding cooperativity mechanism used by transcription factors in absence of strict organisation of their binding motifs, a characteristic feature of most of enhancers.

## Introduction

The binding of transcription factors (TFs) translates the regulatory information contained in cis-regulatory elements (CREs) into gene expression patterns. Upon binding, TFs activate or repress transcription by recruiting protein complexes that modulate the activity of RNA Polymerase II (Pol II) at promoters of genes. The DNA binding domains of TFs recognize 6-25bp DNA sequence motifs with low specificity (Inukai et al., 2017). As a consequence, each individual TF has millions of theoretical recognition sequences in mammalian genomes, few of which are observed to be bound in vivo (Neph et al., 2012). The affinity of a TF for its motif does not explain its genome occupancy, instead combinatorial action of multiple TFs likely shapes the precise control of binding at CREs (Gerstein et al., 2012; Iwafuchi-Doi and Zaret, 2014; Yan et al., 2013). The combinatorial action of TFs is critical for the establishment of gene expression programs during development (Crocker et al., 2015; Farley et al., 2015; Small et al., 1992; Thanos and Maniatis, 1995) and cellular reprogramming (Takahashi and Yamanaka, 2006; Vierbuchen et al., 2010).

Evidence that TFs may collaborate to bind CREs came from the observation that certain key TFs tend to frequently bind the same set of CREs in the genome (Junion et al., 2012; Siersbæk et al., 2014; Tijssen et al., 2011). The interdependency between TFs was further demonstrated by measuring the effects of deletions of individual TFs or their binding motifs on the binding of other TFs in the cluster and/or the activity of the target CREs (Junion et al., 2012; Siersbæk et al., 2014). Complementary evidence for binding dependency comes from comparative genomics showing that binding of a given TF can correlate with changes in the genotype affecting neighbouring binding motifs (He et al., 2011; Kilpinen et al., 2013; Stefflova et al., 2013). These studies have established cooperativity as a prevalent mechanism explaining TF binding at CREs. However, technologies employed in these studies lack the necessary resolution to determine the precise mechanisms of TF cooperativity at specific loci.

Several mechanisms have been described to explain the binding cooperativity of TFs (reviewed in (Deplancke et al., 2016; Inukai et al., 2017; Morgunova and Taipale, 2017; Reiter et al., 2017; Spitz and Furlong, 2012)). Cooperativity was shown to occur through direct protein-protein interactions between TFs. For instance, transcriptional activator Nuclear respiratory factor 1 (NRF1) undergoes homodimerization prior to binding, increasing the affinity of the TF for DNA (Gugneja and Scarpulla, 1997). Other TFs, such as Myc/Max, NFY, and nuclear receptors, undergo physical interactions that combine different TF DNA binding domains to increase their binding affinity and motif specificity (Amoutzias et al., 2008). A systematic study has estimated the existence of >800 interactions between TFs, largely expanding the binding repertoire of the 2000 known human TFs (Ravasi et al., 2010). Cooperativity has also been observed in the absence of direct protein-protein interactions, when the binding of one TF to a locus alters the DNA structure and improves the local affinity for another TF (Jolma et al., 2015). Finally, TF binding cooperativity has been proposed to occur through competition with nucleosomes for DNA occupancy. Wrapping of DNA into nucleosomes forms a physical barrier that TFs must overcome to exert their regulatory function. In this context, DNA binding of a single TF might be insufficient to displace nucleosomes; instead, collective binding of multiple TFs may be required (Mirny, 2010; Polach and Widom, 1995, 1996). Occurrence of this phenomenon in vivo has been demonstrated in principle using artificial systems (Adams and Workman, 1995; Pettersson and Schaffner, 1990; Vashee et al., 1998) but the extent to which this mechanism is used by endogenous TFs remains to be determined.

Mechanistically, an important open question is whether simultaneous co-occupancy of DNA by multiple TFs is required for binding cooperativity. Most TFs have short residence times on DNA (Agarwal et al., 2017; Arnold et al., 2013a; Gebhardt et al., 2013; Sung et al., 2014) and for such TFs, binding cooperativity was already demonstrated to occur in absence of physical co-occupancy (i.e. glucocorticoid receptor (Voss et al., 2011)). Addressing this question requires direct quantification of how frequently two TFs co-occupy the same DNA molecule in vivo. Current methods to measure TF binding, such as Chromatin-Immunoprecipitation (ChIP) are based on enrichment of short DNA fragments bound by the target TFs. These approaches provide precise information on the binding of individual TFs but, they lose information on the co-occurrence of binding at neighbouring sites. To overcome these limitations, we recently developed a Single Molecule Footprinting (SMF) approach for Drosophila melanogaster genomes. We demonstrated its ability to quantify the multiplicity of protein-DNA contacts made by the transcription machinery at the resolution of single DNA molecules genome-wide (Krebs et al., 2017).

Here, we adapt SMF for mammalian genomes and demonstrate that the assay resolves TF binding and nucleosome occupancy at single molecule resolution. We show that SMF allows the simultaneous quantification of multiple TF binding events on single DNA molecules, enabling us to systematically quantify the frequency of co-occupancy for thousands of TF pairs across the genome. Analysis of these binary molecular TF occupancy patterns reveals that simultaneous binding is largely independent of the identity of the factors involved, does not require a strict organization of motifs and is prevalent at regions having high nucleosome occupancy. Reduction of TF concentration using siRNAs indicate that co-occupancy is a mechanism used by TFs to bind their cognate motifs. Altogether, our data comprehensively identify TF interactions at CREs at molecular resolution and show that DNA co-occupancy is a widespread cooperativity mechanism used by TFs to bind DNA and evict nucleosomes.

## Results

### Single Molecule Footprinting of the mouse genome

Accurate quantification of genomic binding events by SMF requires the sequencing of a large number of DNA molecules encompassing the binding regions of the factor of interest. Generation of high coverage SMF datasets is challenging in mammalian genomes, which are a factor of twenty bigger than Drosophila genomes. To overcome this bottleneck, we took advantage of the fact that TF binding is mostly restricted to CREs, which represent only a small fraction of the genome (Stamatoyannopoulos et al., 2012). We employed DNA capture to enrich for CREs prior to sequencing (Figure 1A). We used a library of RNA baits tiling 297,000 regions (2% of the genome), covering a large fraction of promoters, enhancers, and insulators accessible in mouse embryonic stem cells (mESCs) (59.7% of open regions as detected by DNAse-seq).

**Fig. 1.**
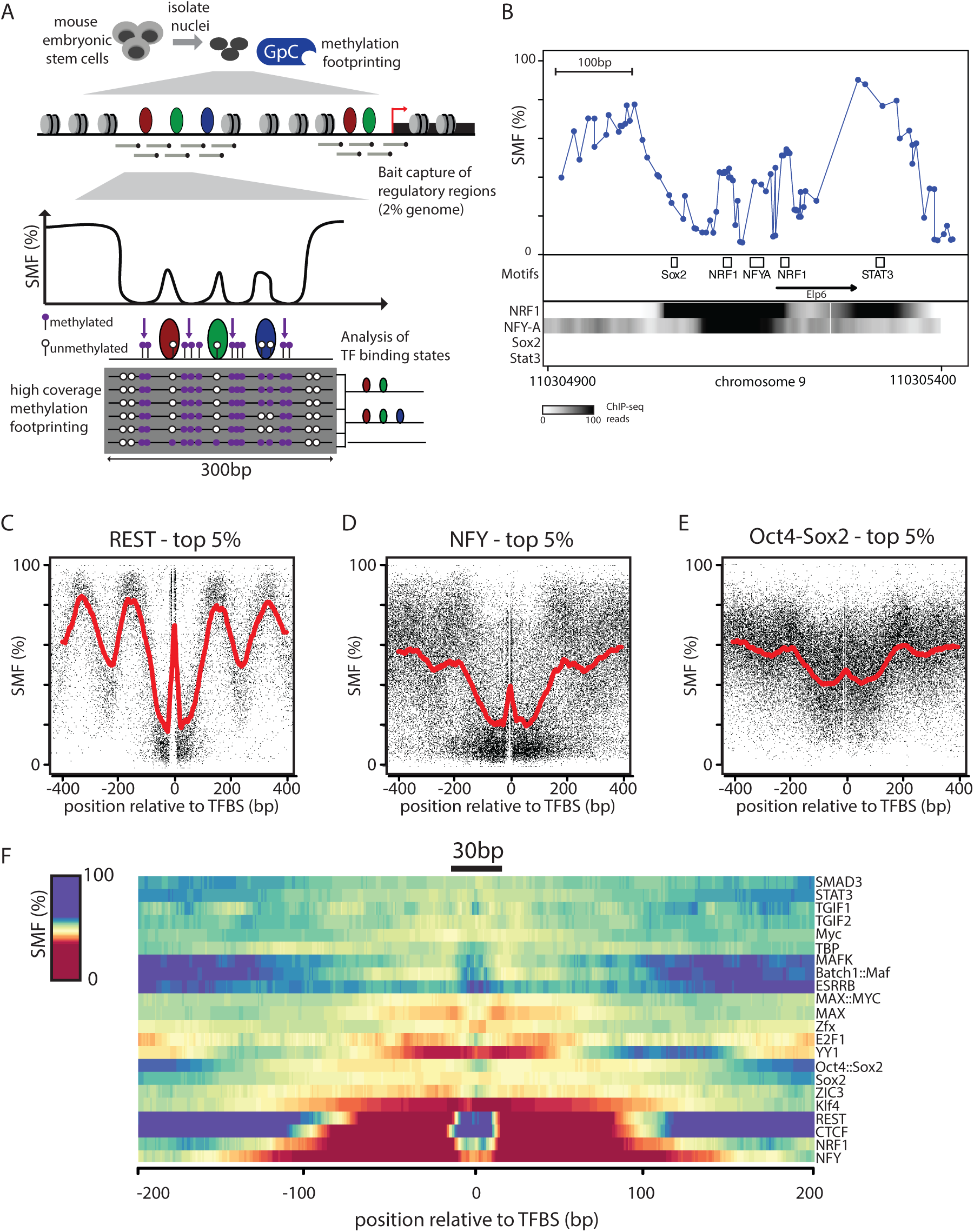
Single Molecule Footprinting detects Transcription Factors in mouse embryonic stem cells. (A) Overview of Single Molecule Footprinting (SMF) for mammalian genomes. Nuclei are isolated and incubated with a recombinant cytosine methyl-transferase that targets GCs (M.CviPI) which are distinct from CGs that are endogenously methylated. To generate data with a genomics coverage compatible with single molecule interpretation, the resulting methylated DNA is subjected to capture using probes targeting 60% of the Cis-Regulatory Elements (CREs) used in mESCs. Methylation is detected by bisulfite sequencing using long reads (300bp), enabling the detection and analysis of multiple footprints on a single molecule. (B) SMF pattern at the active promoter of the Elp6 gene revealing short footprints (<20bp) at binding sites of the activators NRF1 and NFY. Shown is the inverse frequency of methylation (1-methylation (%)) (blue line). Black boxes represent the location of consensus motifs for TFs. Black arrow indicates the transcriptional start site. Read counts for ChIP-seq of the respective TFs are shown as intensity heatmap (datasets as indicated). (C-E) TFs binding creates discrete footprints detectable by SMF. Composite profile of SMF signal at various (C, D, E) bound TF motifs (top 10% of the respective TF ChIP-seq) as indicated. Shown is the footprinting frequency (1-methylation [%]) of individual cytosines (black dots). (F) Identification of the TFs that create footprints in SMF signal. SMF composite profile at the binding sites of 21 TFs depicted as a heatmap. Smoothed average signal around the TFBS of the 10% top sites as defined by the respective TF ChIP-seq.

Unlike Drosophila, mammalian genomes have endogenous DNA methylation which is prevalent in mESCs but almost exclusively limited to CG dinucleotides (Stadler et al., 2011). CG methylation prevents the use of exogenous methyl-transferases targeting this sequence context, reducing the spatial resolution of the assay. We took two complementary approaches to avoid interference between SMF and endogenous DNA methylation. First, we used the cytosine methyltransferase M.CviPI which only methylates GC dinucleotides, therefore compromising on the spatial resolution of the assay (median distance of 14bp). Second, we leveraged the ability of mESCs to proliferate upon genetic ablation of DNA methylation (Tsumura et al., 2006). We used an isogenic mESC line depleted for all three de novo DNA methyl-transferases (DNMT TKO), which shows only small number of localized discrete changes in chromatin accessibility and gene expression (Domcke et al., 2015). This line enables the use of GC as well as CG methyl-transferases for footprinting, significantly increasing the ¬spatial resolution (up to 7bp), which is of considerable utility when analysing discrete TF binding events. However, such analysis is limited to the stem cell state, as DNMT TKO cells fail to differentiate (Sakaue et al., 2010).

We generated high coverage SMF datasets in wild-type and DNMT TKO mESCs with highly reproducible methylation footprints between biological replicates (R>0.90, Supplementary Figure 1 A, B). Moreover, footprinting levels were in close agreement between wild-type and DNMT TKO cells (R=0.90, Supplementary Figure 1C), consistent with previous observations that methylation depletion leads to only discrete changes in the accessibility pattern of mESCs. The high capture efficiency (>70% of reads were within bait regions) achieved coverage levels that allowed data interpretation at the single molecule resolution for 78 807 CREs in mESCs.

### Single molecule detection of TF binding

When inspecting SMF signal at CREs, we frequently observed discrete footprints (<25bp) around TF motifs (as exemplified in the Elp6 promoter, Figure 1B). These motifs are recognized by TFs that are also detectable by ChIP-seq at respective regions (Figure 1B), suggesting that TF binding results in footprints detectable by SMF, consistent with previous observations (Gal-Yam et al., 2006; Kelly et al., 2012; Levo et al., 2017; Oberbeckmann et al., 2019; Shipony et al., 2020). This prompted us to assess whether footprints at TF motifs can be found genome-wide and ask whether the presence of footprints is consistent with orthogonal measures of TF binding. We used a large collection of ChIP-seq datasets of TFs in mESCs (Supplementary Table 1), to identify the subset of motifs that are bound by their TF genome-wide. We plotted the SMF signal around these motifs and found that some factors, such as the transcriptional repressor RE1-Silencing Transcription factor (REST), create short footprints over their bound sites, in contrast with unbound motifs (Supplementary Figure 1F). These footprints are directly flanked by highly accessible regions and larger periodic footprints consistent with nucleosomal phasing (Figure 1C).

We also detected footprints at the binding sites of activators, such as NFY (Figure 1D, Supplementary Figure 1G) and Oct4-Sox2 (Figure 1E, Supplementary Figure 1H). However, the accessibility and the nucleosomal phasing at the flanking regions of these TFs were weaker. A likely explanation for this heterogeneity is the binding of other factors in the vicinity, as these motifs tend to lie within motifs clusters forming CREs. A systematic assessment of footprints at bound motifs revealed that many TFs create footprints detectable by SMF (Figure 1F). These vary in size (15-30bp) and intensity. Thus, methylation footprinting has the sensitivity to discretely quantify TF binding at single molecule resolution.

### Heterogeneity of occupancy at TF binding regions

In higher eukaryotes, TFs bind to a small fraction of their recognition motifs in the genome (Wang et al., 2012). Competition between TFs and nucleosomes is assumed to be a critical determinant of TF binding (Iwafuchi-Doi and Zaret, 2014; Morgunova and Taipale, 2017). ChIP-seq involves selective isolation of DNA bound by a chromatin-associated factor. As a consequence, enrichment does not inform on the proportion of DNA molecules bound by a TF, nor on the level of competition with nucleosomes at a particular target region. To determine the relative fraction of molecules bound by TFs or nucleosomes at individual TF motifs, we adapted the single molecule classification strategy we developed to study the binding of GTFs at core promoters (Krebs et al., 2017). We postulated that combining the accessibility information at an expected TF binding location with flanking genomic locations would enable us to discriminate short footprints, presumably created by TFs, from longer nucleosomal footprints. We created a collection bin at the expected TF binding region and two collections bins on either flank (Supplementary Figure 2A). We collected methylation information in a 15bp window around TF motifs to avoid restricting our analysis to TFs having GCs or CGs in their recognition motifs. For each individual molecule, the algorithm collects binarized methylation within the three bins, creating 8 (23) possible combinations. We further grouped these patterns into three binding states, separating molecules showing a short footprint at the TF motif, fully accessible molecules (no detectible binding at the motif), and molecules showing large nucleosomal footprints (Supplementary Figure 2B).

We applied single molecule quantification of TF binding to the 7383 REST motifs targeted by our capture method. We reproducibly quantified the binding frequency of REST for >77% of the 1357 motifs that contain informative GCs within all three collection bins (Supplementary Figure 2C). The high percentage of recovery confirms the efficiency of our DNA capture strategy and that we reached a sequencing depth allowing single molecule quantification of TF binding.

We observed considerable heterogeneity in the accessibility patterns of bound REST motifs (Figure 2A). For instance, even at a highly bound site only 34% of the DNA molecules showed a short footprint at the motif potentially created by REST (Figure 2A, left panel). Others were accessible (17%) or harboured larger footprints compatible with nucleosomal occupancy (49%). If these short footprints are created by REST, genetic deletion of the TF should abolish them. We compared REST single molecule profiles with those obtained in cells genetically depleted for REST (REST-KO) at the same locus (Chen et al., 1998) (Supplementary Figure 2D). In the absence of REST, the discrete footprint at the binding site disappeared and almost all molecules showed the large footprints assigned to nucleosomes (92%) (Figure 2A, right panel). A complete loss of presumptive TF binding was observed for all 16 REST binding sites tested (Figure 2B), indicating that these are created by REST binding.

**Fig. 2.**
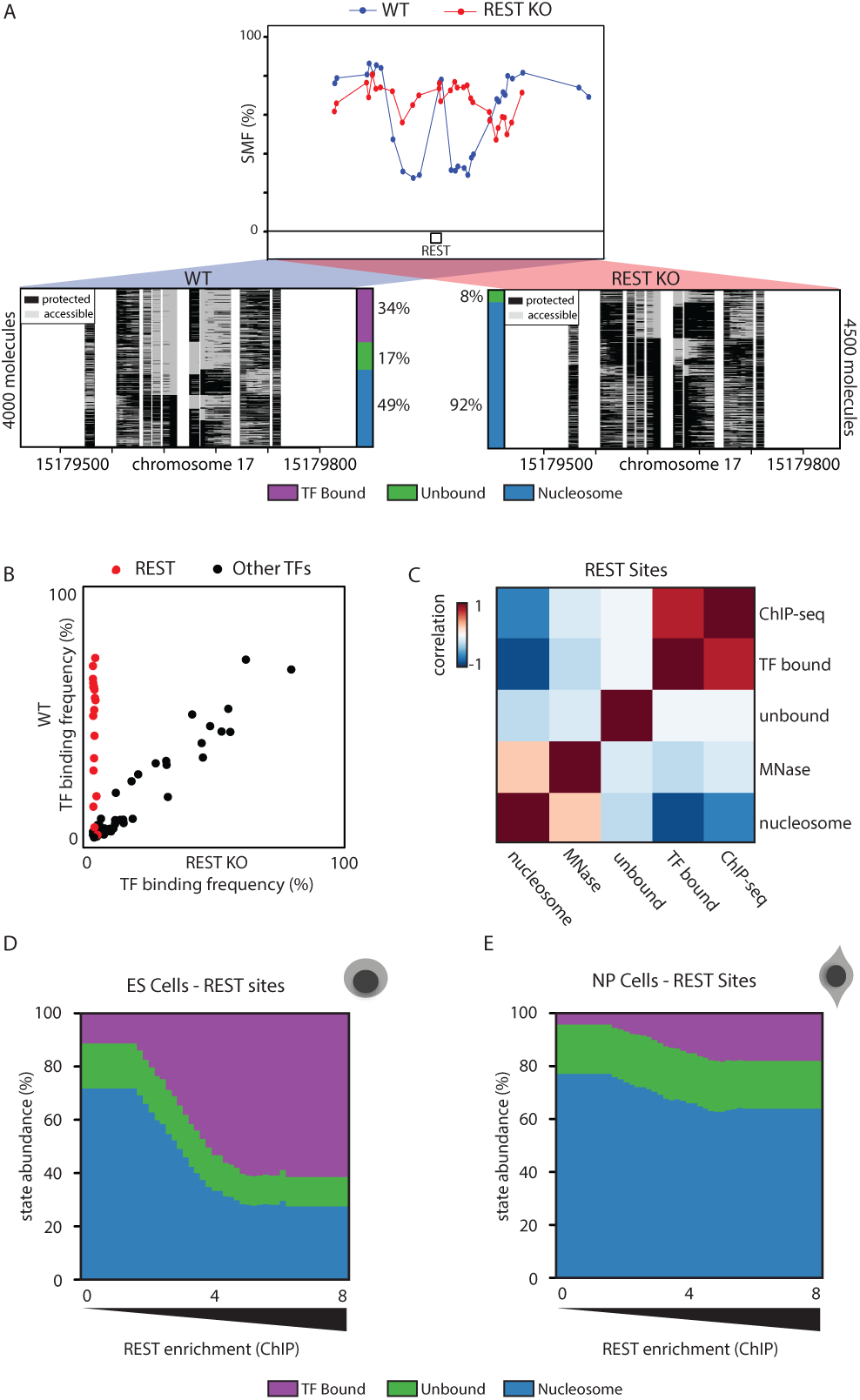
Quantification of TF binding frequency at single DNA molecule resolution. (A) Single-locus examples of a REST bound region in wild type (WT, blue line and dots) versus REST knock out (REST-KO, red line and dots) mESCs. Shown are average methylation levels (top panel, blue dots connected by a blue line) and single-molecule stacks (bottom panels) measured by targeted amplicon bisulfite sequencing, sorted into three states using the classification algorithm (methylated Cs, accessible, light gray; unmethylated Cs, protected, black). The vertical side bar depicts the frequency of each state. Color legend for the states is given on the bottom of the panel (purple: TF bound; green: accessible; blue: nucleosome occupied). The percentages of molecules in each state are indicated on the right side of the plot. (B) Scatter plot comparing TF binding frequencies in WT and REST KO mESC at 16 REST motifs (red) and motifs of other TFs (black) covered by 96 amplicons using targeted SMF. (C) Global relationship between state frequencies and independent bulk measurements of TF and nucleosome occupancies at REST motifs, depicted as a heatmap of similarity (Pearson correlation). States separate in two groups that either correlate with occupancy by the TF (ChIP-seq) or nucleosomes (MNase-seq) illustrating accurate state quantification. (D-E) Distribution of state frequencies in mESC (D) and neuronal progenitors (NP) (E) as a function of REST occupancy as determined by ChIP-seq. Cumulative bar plot depicting the distribution of state frequencies. TFBS were binned based on REST enrichment in mESCs (log2 ChIP-seq), and the median frequency of each state was calculated within each bin. The frequency of each state is color coded as in Figure 2A. TF occupancy changes across loci and between cell types are tightly coupled with nucleosome occupancy.

If SMF accurately quantifies TF binding and nucleosome occupancy, then the frequency of the TF footprints should scale with TF and nucleosome occupancy as determined by bulk assay. Comparison with existing REST ChIP-seq and MNase-seq data at REST binding sites revealed that the SMF frequency of the TF bound state was strongly correlated with enrichments as measured by ChIP-seq (R=0.74, Figure 2C). Conversely, frequency of the nucleosome occupied state showed a correlation with nucleosome occupancy at the TF binding site as measured by MNase-seq (R=0.3, Figure 2C). We conclude that SMF simultaneously quantifies TF and nucleosome occupancy, revealing the heterogeneity of binding patterns at CREs.

### TF/nucleosome competition at REST binding sites

The heterogeneity observed in SMF patterns suggests that within a cell population, most REST motifs can either be occupied by REST or by nucleosomes, indicating a competition between these two states. If so, variations in TF binding and nucleosome occupancy levels should be tightly and inversely coupled when compared across these regions. To test this idea, we analysed the relationship between TF and nucleosome occupancy at motifs, as a function of REST binding intensity (Figure 2D). The frequency of TF bound molecules grew with increasing ChIP-seq enrichment, with up to 60% occupancy for the top TF bound sites. Increases in TF occupancy were accompanied by a proportional decrease in nucleosome occupancy (Figure 2D). Overall, nucleosomes occupy 20-60% of the REST bound sites (Figure 2D), implying that at any given time only a fraction of the cells undergo TF binding at a particular binding site.

The tight coupling between the states suggests there is competition between nucleosome occupancy and REST binding at these sites. To further test this possibility, we investigated how binding states redistribute upon perturbation of REST expression levels. During the early stages of neuronal differentiation, REST expression levels are reduced by 4fold (Supplementary Figure 2E) without significant redistribution of REST target sites (Arnold et al., 2013b). We generated a SMF dataset in in vitro derived neuronal progenitors (NPs) (Bibel et al., 2004) (Supplementary Figure 1D, E). Upon reduction of REST expression, we observed a global decrease of 4-fold in REST occupancy at its motifs (Figure 2E), indicating that REST abundance correlates with its binding frequency in the cell population. We also observed a concomitant increase in nucleosome occupancy at REST sites (Figure 2E), whereas there was very little effect on CTCF binding frequencies at CTCF binding regions in NPs (Supplementary Figure 2G, H). Thus, the heterogeneous patterns at REST binding regions are the result of competition between the TF and nucleosomes for DNA occupancy.

### TFs have diverse nucleosome remodelling patterns

A small number of TFs bind their target sites in isolation (i.e. REST, CTCF), but most bind within larger clusters of TF motifs at enhancers and promoters. To understand how this functional diversity influences the single molecule occupancy of TFs, and how it relates to alterations of chromatin structure, we classified the footprint patterns around all binding sites for which binding motifs are known (500 from JASPAR (Mathelier et al., 2015)). We subsequently selected the motifs for which ChIP-seq data were available in mESCs (Supplementary Table 1), and obtained genome-wide TF binding frequencies for 22 TFs that were highly correlated between biological replicates (Supplementary Figure 3). For this TF collection, we asked how the frequency of unbound, TF bound, or nucleosome occupied DNA molecules quantitatively compare with TF enrichment as measured by ChIP-seq (Figure 3A). We observed that the insulator CTCF has very similar binding characteristics to REST (Figure 3A; category 1), where molecular occupancy by the TF correlates closely with ChIP-seq enrichment (Figure 3A). For these two factors, we observe that DNA molecules are bound by either the TF or nucleosomes with a constant but small (10%) fraction of unbound DNA molecules (Figure 3B, E).

**Fig. 3.**
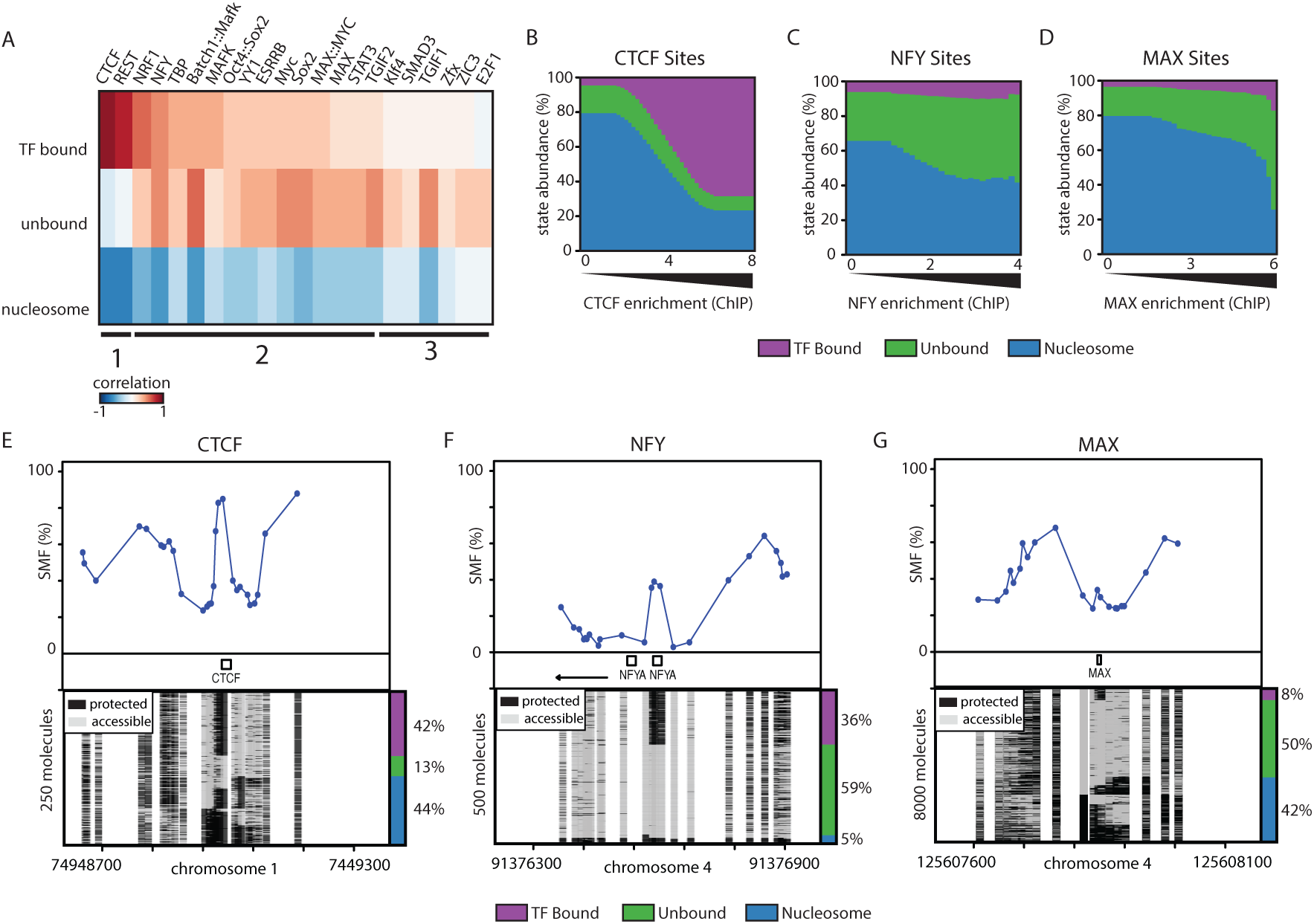
Identification of the nucleosome remodeling strategies used by TFs. (A) Global relationship between SMF state frequencies and TF occupancy as determined by ChIP-seq at TF motifs. Heatmap of similarity (Pearson correlation) between the frequency of states (y-axis label) at specific motifs and ChIP-seq for the matching TF (x-axis label). Columns are ranked by similarity between the TF bound state and ChIP-seq. The side bar indicates groups of TFs sharing similar profiles. (B-D) Distribution of state frequencies as a function of TF occupancy as determined by ChIP-seq for (B) CTCF, (C) NFY, (D) MAX. Cumulative bar plot depicting the distribution of state frequencies. TF motifs were binned based on their respective ChIP-seq enrichment in mESCs (log2 ChIP-seq), and the median frequency of each state was calculated within each bin. The frequency of each state is color coded as in Figure 2A, color legend for the states is given on the bottom of the panels. (E-G) Single-locus examples of a (E) CTCF, (F) NFY, (G) MAX bound region mESCs. Shown are average methylation levels (top panel, blue dots connected by a blue line) and single-molecule stacks measured by targeted amplicon bisulfite sequencing, sorted into three states using the classification algorithm (methylated Cs, accessible, light gray; unmethylated Cs, protected, black). The vertical side bar depicts the frequency of each state. A color legend for the states is given on the bottom of the panels. The percentages of molecules in each state are indicated on the right side of the plot.

For a majority of the other tested TFs, the occurrence of TF footprints also scaled with ChIP-seq enrichments, but reached lower maximal frequencies at the highest bound motifs (Figure 3A, C, D). In this category, a large fraction of DNA molecules was unbound (Figure 3A; category 2), as exemplified for the activators NFY or MAX (Figure 3C-D, F-G). For a smaller set of factors, we observed a good correlation between unbound DNA molecules and ChIP-seq enrichments with very infrequent TF-bound DNA molecules (Figure 3A; category 3). The different behaviours of the categories may be due to TF residence times on DNA (Voss and Hager, 2014), or TF concentration in the nucleus. For instance, TFs in categories 2 and 3 are activators that were found to have shorter residence times than CTCF (Agarwal et al., 2017; Voss and Hager, 2014).

The degree of anti-correlation between the nucleosome-occupied fraction and ChIP-seq enrichments is lower for categories 2 and 3, compared to category 1 (Figure 3A). For factors in categories 2 and 3, a significant fraction of DNA molecules is unbound, even at low ChIP-seq enrichments (exemplified in Figure 3C, F). The frequency of these unbound DNA molecules increases at higher ChIP-seq enrichment. Together, this suggests that nucleosome remodelling can only be partially explained by the binding of individual factors at these regions, unlike our observations for REST (Figure 2D, Figure 3B). This is consistent with the idea that many of these binding events occur within clusters of motifs that may collectively contribute to nucleosome remodelling. We conclude that many of the tested TFs create footprints quantifiable at the single molecule level, but that the range of binding frequencies varies greatly from one TF to the other. Finally, our data suggests that combinatorial mechanisms underlie nucleosome eviction for most of the tested TFs.

### Quantification of the molecular co-occupancy of TFs

Having established the ability of SMF to quantify binding of individual TFs on single DNA molecules, we developed a strategy to quantify their degree of molecular co-occupancy (see methods section, Supplementary Figure 4A). We took advantage of the dual enzyme footprinting dataset generated in DNMT TKO cells, which enables quantification of approximately five times more binding events than experiments in WT cells (Supplementary Figure 4B). We did not observe major differences in TF binding frequencies when comparing DNMT TKO cells to WT cells (R=0.94, Supplementary Figure 4C). In our analysis, we distinguished co-occupancy at dimeric motifs (Figure 4A) from TFs occupying distinct motifs lying in the vicinity of each other (15-140bp, Figure 4B). When applying this strategy to the TFs creating quantifiable footprints by SMF (Figure 3A), we obtained reproducible high confidence co-occupancy measurements for 1238 TF dimers and 381 TF pairs (Supplementary Figure 4B, C).

**Fig. 4.**
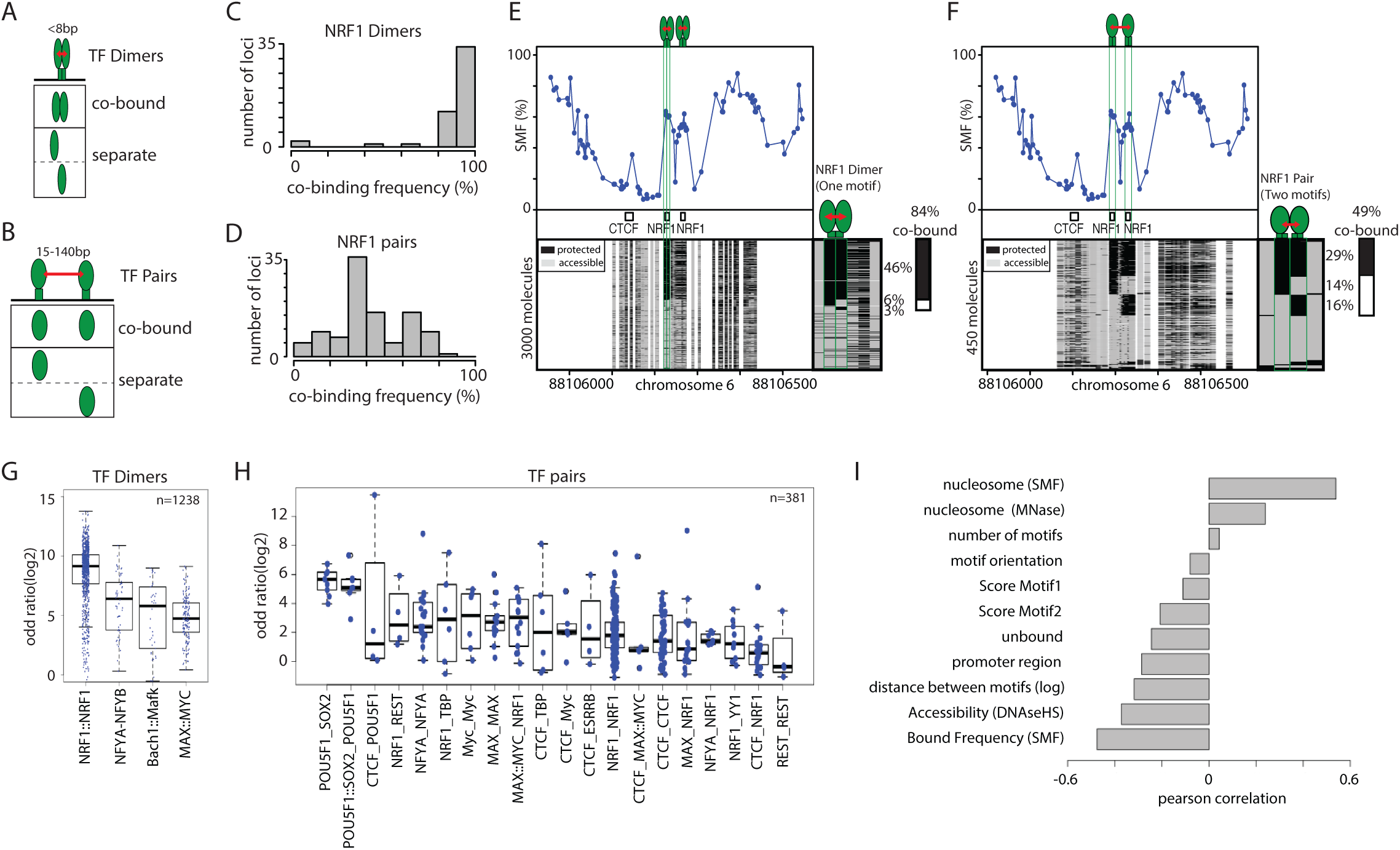
High TF co-occupancy does not rely on a precise motif arrangement. (A) Footprints for dimeric TFs occur nearby (<8bp) on two halves of the same recognition motif. Schematic representation of the theoretical binding states expected to occur at such motifs, bound by TF dimers. (B) Footprints for neighboring binding sites occurs at separate recognition motifs within the same CRE (<140bp). Schematic representation of the theoretical binding states expected to occur at a pair of neighboring motifs. (C) Histogram depicting the percentage of dimeric co-occupancy at NRF1 motifs. (D) Histogram depicting the percentage of co-occupancy at neighboring pairs of NRF1 motifs. (E, F) Single-locus example of a region bound by NRF1 at two neighboring recognition motifs. Analysis of the degree of TF co-occupancy (E) within the NRF1 dimeric motif and (F) between neighboring motif pairs. Shown are average methylation levels (top panel, blue dots connected by a blue line) and single-molecule stacks measured by targeted amplicon bisulfite sequencing (bottom left), sorted into four states using the co-occupancy classification algorithm (methylated Cs, accessible, light gray; unmethylated Cs, protected, black). The states heatmap (bottom right, binarized heatmap) depicts the occupancy states for each of the protein analyzed in the pairs. The percentages of molecules where individual or co-binding are observed is indicated on the right side of the plot. The fraction of co-bound molecules within all bound molecules is indicated on top of a side bar depicting the co-bound frequency. (G-H) Half motifs are frequently co-bound by TF dimers while motif pairs show highly variable degree of co-occupancy. Degree of co-occupancy observed at (G) half-motifs bound by TF dimers, and (H) at neighboring motifs bound by TF pairs. Boxplot summarizing the odds ratios of a Fischer’s exact test for each pair of TFs analyzed. Blue dots represent individual TF pairs. (I) Analysis of the determinants of TF co-occupancy. Barplot depicting the Pearson correlation coefficient between the degree of co-occupancy of TF pairs and various genomic features (y axis label).

Many active promoters harbour clustered motifs for the transcriptional activator NRF1, which binds a tandem repeat recognition sequence as a homodimer. We analysed the frequency of co-occupancy of NRF1 monomers when binding the two halves of the NRF1 motif (Figure 4A), and compared these with the co-occupancy of full NRF1 motifs located within the same CRE (Figure 4B). We observed nearly systematic co-occurrence of footprints at the NRF1 half-sites (>80% of the TF bound molecules, Figure 4C). In contrast, the frequencies of co-occupancy observed between neighbouring NRF1 motifs ranged from very low (<20%) to levels comparable to those between dimeric half-sites (>80%) (Figure 4D). This is also evident when analysing co-occupancy within half-sites (85%, Figure 4E) and between two NRF1 binding sites (48%, Figure 4F) of a single locus. Together, these data provide evidence that NRF1 binds as an obligatory homodimer in vivo and suggests that binding dependency between neighbouring NRF1 binding sites varies substantially from one CRE to another.

SMF is performed on permeabilized nuclei, in the absence of protein-DNA cross-linking. The residence times for TFs in living cells range from a few seconds to a couple of minutes, which are much shorter than the time nuclei are the incubated with methyltransferases in vitro. Thus, it is possible that the co-occupancy patterns detected by SMF could reflect TF retention on chromatin in vitro, rather than binding dependencies between TFs occurring in vivo. To address this question, we developed an independent SMF protocol where we fixed protein-DNA interactions in vivo using formaldehyde prior to methylation footprinting (X-link SMF, see methods section). We observed minimal differences in TF binding frequencies when comparing SMF data between native and crosslinked conditions (Supplementary Figure 4D). Moreover, TF co-occupancy profiles of individual loci were very similar at the single molecule level (Supplementary Figure 4E, F). Together, these findings show that SMF reflects the binding behaviour of TFs in vivo and that co-binding frequencies between TFs is highly variable at CREs.

### Co-occupancy is largely independent of TF identity

To test if co-occupancy patterns are specific to the type of TFs involved, we compared the results obtained for NRF1 with those for other pairs of TF dimers or neighbouring binding motifs. For each binding event, we tested whether the observed TF co-occupancy exceed the one expected by chance. Consistent with our observations for NRF1, we observed that co-occupancy at the half-sites of dimeric motifs is often higher than expected by chance (Figure 4G). However, none of the tested cases showed a systematic co-occupancy that would suggest that obligatory dimerization comparable with what was observed for NRF1 dimers (Figure 4G). For example, we observed that the tandem motif bound by the transcriptional activators Myc and Max showed varying degrees of co-occupancy across the genome (Figure 4G) and we could identify Myc-Max bound regions where >80% is monomeric (Supplementary Figure 4F) This is in agreement with in vitro evidence that Myc-Max dimer formation preferentially occurs through a sequential monomer binding pathway (Kohler et al., 2002). We observe that co-occupancy at most dimeric motifs is higher than random, but not systematic. Our data argues that dimerization is not a prerequisite for binding for most of the tested dimeric factors.

The degree of co-occupancy between pairs of TFs that bind distinct motifs (Figure 4H) is on average lower than the one we observed for TF dimers (Figure 4G). When grouping the data according to the identity of the TFs involved, certain pair types tend to have higher co-occupancy than others, for example, the Pou5f1-Sox2 pairs (Figure 4H). However, we also observed a broad distribution of co-occupancies across pairs formed by the same TFs at different loci. For instance, we observed NRF1-NRF1 pairs with over 80% of co-occupancy at certain sites, but also lower than 20% cooccupancy at other sites (Figure 4D, H). Thus, certain TF combinations tend to co-occupy DNA more frequently than others. However, most variation in TF pairs co-occupancy cannot be explained by the identity of the TFs involved.

### Distance between motifs contributes to high co-occupancy

Most of the tested pairs of TFs co-occupy DNA at high frequency at a subset of their binding loci. Thus, increased co-occupancy is unlikely to require specific proteinprotein interaction domains. To identify potential determinants of the variation observed in TF co-occupancy patterns, we tested how much motif organisation (i.e. number of motifs, their score, distance and orientation) and the binding frequencies of TFs and nucleosomes predict the observed levels of co-occupancy (Figure 4I). Several of the tested parameters show a significant correlation with the degree of TF cooccupancy, including the distance between the binding sites, nucleosome occupancy at the binding region, and TF occupancy level (Figure 4I). However, high co-occupancy does not appear to require precise orientation of the motifs (Figure 4I).

The degree of co-occupancy between TFs decreases as a function of the distance between their binding sites (Figure 5A). Nearby binding events (within 20bp) show greater co-occupancy (Figure 5A). The frequencies of co-occupancy exponentially decrease with increasing distance between the binding sites, with close to random distribution of binding for sites located at greater than 70bp apart (Figure 5A). Similar correlations were seen when considering TF identity (Figure 5B). A large majority of NRF1-NRF1 pairs were frequently co-bound when located at <40bp distance (Figure 5E), whereas co-occupancy was significantly reduced when the sites were located further apart (Figure 5F). However, while distance is an important determinant of co-occupancy, it only partially explains the co-occupancy levels observed at intermediate distances (20-70bp, Figure 5A, B). In summary, co-occupancy between TFs increases with proximity but this does not rely on precise spacing or orientation of the motifs.

**Fig. 5.**
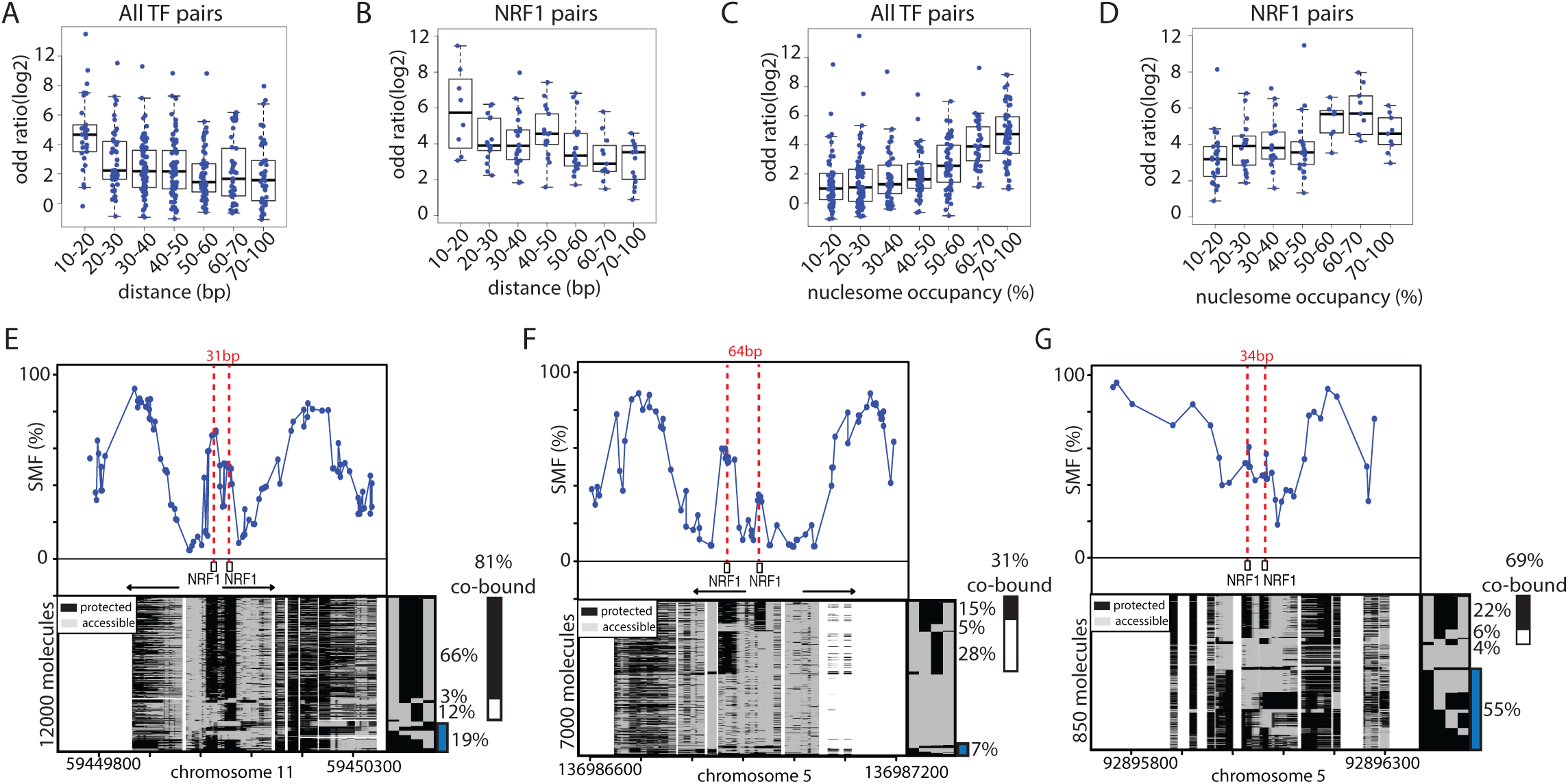
Increased TF co-occupancy occurs at nucleosome occupied regions. (A-B) Co-occupancy at pairs of motifs decrease as a function of genomic distance. Boxplot depicting the degree of co-occupancy as a function of the genomic distance between the TF motifs for all (A) analyzed TFs (B) pairs of NRF1 motifs. Shown are odds ratios of a Fisher exact test. (C-D) Co-occupancy between pairs of motifs is increased at regions having high nucleosomal occupancy. Boxplot depicting the degree of co-occupancy as a function of the fractions of molecules occupied by nucleosomes for all (A) analyzed TFs and (B) pairs of NRF1 sites. (E-F) Comparison of single loci harboring (E) nearby or (F) distant tandem NRF1 motifs. Same representation as in Figure 4E. Blue bar depicts the fraction of nucleosome occupied molecules (G) Example of a NRF1 bound locus having high nucleosome occupancy. Same representation as in Figure 5E.

### TF co-occupancy is high at nucleosome occupied regions

Another predictor of TF co-occupancy is the fraction of DNA molecules competitively occupied by nucleosomes at the same locus (Figure 4I). We observed an increase in TF co-occupancy for regions where more than half of the DNA molecules are occupied by nucleosomes (Figure 5C). While we detect this increase regardless of TF identity (Figure 5C), it is also observed when selecting specific TF pairs (Figure 5D). This effect is evident when comparing two loci bound by the same combination of TFs but having different degrees of occupancy by nucleosomes. A locus with low nucleosome occupancy (7% Figure 5F) has lower TF co-occupancy than a locus with high nucleosome occupancy (55% Figure 5G). Regions of high nucleosome occupancy are likely to be regions where TFs and nucleosomes are actively competing for DNA occupancy, leading to an equilibrium between the two states. In this case, binding of individual TFs would be insufficient to overcome the energetic costs of nucleosome eviction, leading to a requirement for binding cooperativity between TFs at the locus.

### TF co-occupancy identifies cooperativity between TFs

Our analysis of steady-state level of occupancy by TFs and nucleosomes suggests a model in which cooperativity involving competition with nucleosomes mechanistically occurs through physical co-occupancy of the DNA molecules by TFs. If correct, perturbation of the binding of TFs with high frequency of co-occupancy should impact the binding of their partners and nucleosome occupancy at those loci. To test this hypothesis, we performed siRNA knock-down (KD) of NRF1. NRF1 has a broad range of co-occupancy frequencies and is involved in heterologous pairs with many other TFs (Figure 4H). Upon KD, we observed a reduction in NRF1 protein levels (Supplementary Figure 5A) and an average decrease of about one third of NRF1 occupancy at all its binding sites (intercept=0,68, Figure 6A), in contrast with the binding frequencies of other TFs, which are generally unaffected (Figure 6A). Concomitant with the loss in NRF1 binding, we observed a proportional gain of nucleosome-occupied DNA molecules at the NRF1 bound sites (Supplementary Figure 5B).

**Fig. 6.**
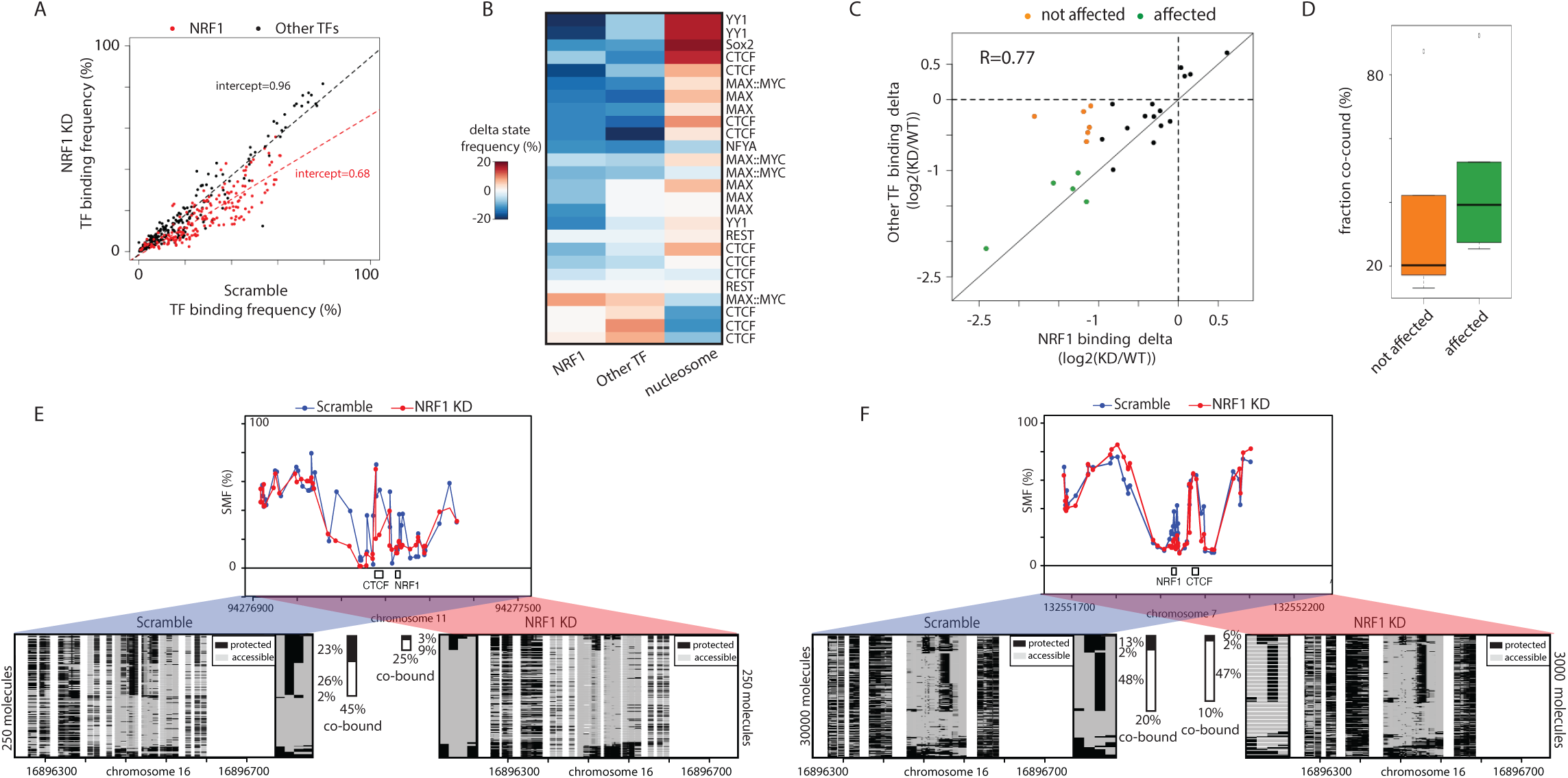
TF co-occupancy is a mechanism of TF binding cooperativity. (A) Specific reduction of TF footprints at NRF1 motifs upon NRF1 knock down (KD). Scatterplot depicting the binding frequency of TFs upon NRF1 KD. Binding frequency at NRF1 motifs is consistently decreased (red dots) while other TFs are mostly unaffected (black dots). Dotted lines represent a linear regression fitted to NRF1 (red) and the other TFs (black). A proportional loss of 30% is observed at NRF1 binding motifs. (B) Decrease in NRF1 binding affects binding of heterologous factors at neighboring motifs. Heatmap depicting the changes in binding frequencies for NRF1-containing heterologous motif pairs. Shown is the difference in TF binding frequency and nucleosome occupancy between WT and NRF1 KD. The identity of the second TF is indicated on the row labels of the heatmap. The rows were grouped using k-means clustering. (C) Binding changes are correlated at most NRF1-containing heterologous TF pairs. Scatterplot comparing the loss at NRF1 binding motifs with the one at neighboring heterologous factors. A fraction of the TFs involved in heterologous pairs with NRF1 have correlated reduction of their occupancy upon NRF1 KD (green dots), while other are not affected (orange dots). (D) Binding frequency is decreased upon KD for TFs having high co-occupancy with NRF1. Boxplot depicting the frequency of co-occupancy for TFs that are not affected (orange) or strongly reduced (green) by NRF1 KD. Categories are similar to Figure 6C. (E-F) Single-locus examples of CTCF bound regions where (E) high or (F) low binding cooperativity with NRF1 is observed. Shown is the average SMF signal in mESCs treated with scramble (top panel, blue dots connected by a blue line) or NRF1 siRNA (red dots connected by a red line). Same representation as in Figure 4E.

We observed that upon KD, a majority of TF binding events involved in a heterologous pair with NRF1 showed reduced TF binding (Figure 6B). Importantly, concomitant to this loss we observed an increase in nucleosome occupancy at these sites (Figure 6B). NRF1 KD affected diverse transcriptional activators but also a subset of the binding sites of the insulator CTCF. We observed a good correlation between loss of NRF1 binding and reduction in binding of the heterologous TF (Figure 6C), suggesting that this reduction is a direct effect of the decrease in NRF1 binding. However, we could identify a set of TF binding events that were less affected by the reduction of NRF1 binding in their vicinity (Figure 6C – orange dots). These less affected TF binding events tend to have a lower frequency of co-occupancy with NRF1 than those showing a substantial reduction upon NRF1 KD (Figure 6D). This is further evident when comparing the important decrease in CTCF binding frequency at a locus where NRF1 co-occupies 46% of its bound molecules (Figure 6E) with the limited changes observed at a locus where it only co-occupies 22% CTCF’s bound molecules (Figure 6E). Thus, high TF co-occupancy is consistent with cooperativity between NRF1 and its partners at these sites. Together our data suggest a cooperativity model in which increased TF co-occupancy is required to open chromatin at sites of high nucleosome occupancy.

## Discussion

Here, we demonstrate the applicability of single molecule footprinting (SMF) to quantify the binding of TFs and nucleosomes at mouse CREs. We show that the resolution of SMF is sufficient to simultaneously detect the binding of multiple TFs on single DNA molecules. We use this property to systematically quantify the degree of co-occupancy of neighbouring TF binding events. We find widespread evidence of co-occupancy for most of the TFs types tested and demonstrate that high co-occupancy identifies binding cooperativity between TFs. Our data identifies TF co-occupancy as an important mechanism used by TFs to evict nucleosomes in order to access their binding sites.

We detected quantifiable footprints for more than half of the TFs we tested in mESCs. For these TFs, the frequency of TF footprints scales with orthogonal measures of TF binding using ChIP-seq. Differences in in vivo footprint patterns between TFs has been previously reported for other DNA footprinting technologies, such as ATAC-seq or DNase-seq (Karabacak Calviello et al., 2019; Neph et al., 2012) and may result from differences in the residence time of TFs on DNA (Agarwal et al., 2017; Arnold et al., 2013a; Gebhardt et al., 2013; Sung et al., 2014). We did not observe major changes in footprint patterns when performing SMF on chromatin preparations where binding events are integrated over several minutes by formaldehyde crosslinking. This is in agreement with previous observations showing that not all yeast TFs create footprints using methylation footprinting under crosslinking conditions (Levo et al., 2017), and is consistent with the subtle differences observed in DNase-seq or ATAC-seq footprints when performed on crosslinked material (Oh et al., 2019). In SMF data, accessibility can only be probed in specific dinucleotide contexts occurring at every 7-14bp. To avoid restricting our analysis to TFs having GC or CGs in their binding motifs, our analysis uses a binning strategy that extends beyond the TF motif. Nevertheless, with these limitations in the spatial resolution of the data, only a fraction of the binding sites is quantifiable at single molecule resolution (i.e. 28% for REST). This could possibly be improved in the future by using methyltransferases targeting other nucleotide contexts (Shipony et al., 2020).

Simultaneous measure of TF binding and nucleosome occupancy revealed the various strategies adopted by TFs to evict nucleosomes. For instance, the zinc-finger domain containing TFs CTCF and REST show a tight coupling between TF binding and nucleosome occupancy. Despite the similarity in their relationship to chromatin remodelling, these TFs depend on different remodelling complexes to evict nucleosomes (Barisic et al., 2019). Another common characteristic of these TFs is that they tend to bind in isolation in the genome. Therefore, chromatin remodelling is likely to depend on the sole binding of these TFs at their target motifs. In contrast, the target motifs for most TFs are found within clusters (Vierstra et al., 2020). We observe that TFs typically binding such clusters occupy only a fraction of the potential target DNA molecules, suggesting that chromatin remodelling at these sites is the result of the collective action of multiple TFs. Thus, typical enhancers, which consist of clusters of TF binding sites, are largely depleted for nucleosomes, and are bound by TFs in heterogenous patterns across the cellular populations.

The unique ability of SMF to detect the binding of multiple TFs on single DNA molecules enabled to determine their frequency of co-occupancy at CREs. As expected, we found that the frequency of co-occupancy is increased at known dimeric sites where protein-protein interactions are expected to occur. However, for most of the known TF dimers tested, dimerization is not obligatory for in vivo binding, suggesting that sequential monomer binding is the binding mode used by many TF dimers. Interestingly we also observed comparably high frequency of co-occupancy for TFs that are unlikely to physically interact when these are binding at regions having high nucleosome occupancy. This shows that protein-protein interactions are not mandatory for high TF co-occupancy, but are likely to contribute to the stabilisation of TF bound complexes in vivo. These observations make predictions on how TF binding motifs are organised at CREs (Spitz and Furlong, 2012) and has implications for the models of enhancer activity (Arnosti and Kulkarni, 2005; Spitz and Furlong, 2012). Our data argue that strict motif organisation is dispensable for TF cooperativity at most CREs but that protein-protein interactions lead to stabilisation of TF binding at dimeric motifs. This agrees with a ‘billboard’ model for CREs, where motif organisation is generally flexible with the occurrence of enhanced cooperativity modules requiring stricter motif organisation.

Our data suggests that TF co-occupancy is largely dictated by the equilibrium state between TF binding and nucleosome occupancy at CREs. This phenomenon is independent of a precise arrangement of binding sites, but is exponentially reduced with genomic distance between the motifs. These observations are best explained by a model of nucleosome-mediated cooperativity for TF binding as formulated by (Mirny, 2010). Thus, we provide formal evidence that nucleosome-mediated cooperativity is widely used by TFs to access DNA at endogenous regulatory regions. Our data further suggest that nucleosome eviction is a direct consequence of simultaneous binding of TFs rather than their additive action upon dynamic binding, refining the mechanistic model of TF cooperativity driven by competition with nucleosomes.

## Methods

### Experimental model and subject details

Wild-type ES 159 and DNA methylation-null ES (DNMT TKO) cells were grown on gelatin-coated 10cm dishes, using Dulbecco’s Modified Eagle Medium (DMEM), supplemented with 15% FBS, LIF, 2-Mercaptoethanol, 1mM L-Glutamine and 1x non-essential amino acids (ThermoFisher 11140050) with regular splitting of cells.

### Single Molecule Footprinting

Footprinting protocol was adapted from Krebs et al 2017 and optimized for mESC. Per reaction, 250 10^3 mESC (ES 159 or DNMT TKO) were trypsinized, washed in ice-cold PBS and re-suspended in icecold hypertonic lysis buffer (10mM Tris (pH = 7.4), 10mM NaCl, 3mM Mgcl2, 0.1mM EDTA, 0.5% NP40). Cells were incubated 10min on ice, releasing nuclei and span down, pelleting the nuclei. Nuclei were washed with SMF Wash Buffer (10mM Tris (pH = 7.4), 10mM NaCl, 3mM Mgcl2, 0.1mM EDTA) and re-suspended in 1x M.CviPI reaction buffer (50mM Tris (pH 8.5), 50mM NaCl, 10mM DTT). The nuclei were then incubated with 200U of M.CviPI (NEBM0227L) at 37°C for 7.5 min (in presence of 0.6mM SAM, and 300mM Sucrose). The reaction was supplemented with 100U of M.CviPI and 128pmol of SAM before a second incubation round of 37°C for 7.5 min. For ES samples, reactions were stopped at this point by adding a SDS containing buffer (20mM Tris, 600mM NaCl, 1%SDS 10mM EDTA). For DNMT TKO samples, where dual enzyme footprinting was applied, 10mM MgCl2, 128pmol of SAM and 60U of M.SssI (NEB- M0226L) were added for a third incubation round of 37°C for 7.5 min followed by stoppage of the reaction by adding the same SDS containing buffer. For all samples, material was digested with Proteinase K (200 mg/ml) overnight at 55°C, followed by phenolchloroform purification of DNA, which was resuspended in water and treated with RNase A at for 1h 60C and quantified using Qubit 1x dsDNA High Sensitivity kit.

### Bait-capture of footprinted DNA

Genome-wide data were obtained using Agilent SureSelectXT Mouse Methyl-Seq Kit. The library preparations for the genome-wide data were performed according to the SureSelect XT Mouse Methyl-Seq Kit Enrichment System for Illumina Multiplexed Sequencing Library protocol (Agilent Technologies, Santa Clara CA, Version E0, April 2018). A total of 3 µg of footprinted DNA was used as input for bait capture, according to the company’s specifications. (5190-4836). DNA was first sonicated using a Covaris S220 sonicator (Covaris, Woburn, MA) (duty factor 10%, intensity 4 and 200 cycles/burst for 100 sec duration time) to obtain products of 200-300 bp. DNA was then end-repaired, A-tailed and ligated with methylated adapters to create a pre-capture DNA library. Adapter-ligated libraries were purified using (0.65X) AMPure XP beads (Beckman Coulter, Inc., Brea, CA, USA) then quality and quantity of libraries were determined by bioanalyzer using DNA high sensitivity chip (Agilent). Next, 350 ng of each library was hybridized with the SureSelect Mouse methyl-seq capture library at 65 °C for 16 h. Hybridized products were purified by capture with Dynabeads MyOne Streptavidin T1 magnetic beads (ThermoFisher Scientific, Waltham, MA)and then subjected to bisulfite conversion using EZ DNA Methylation- Gold Kit (Zymo Research D5005)kit according to manufacturer’s protocol. As described in manufacturer’s protocol (Agilent SureSelectXT Mouse Methyl-Seq Kit), bisulfite converted libraries were PCR-amplified for 8 cycles with supplied universal primers and purified using AMPure XP beads. Captured libraries were indexed by PCR for another 6 cycles, using supplied indexes for downstream multiplexing. Quality of libraries were ensured with an Agilent Bioanalyzer using DNA High Sensitivity chip prior to pooling for sequencing. Supplied primers and recommended amplification parameters of the manufacturer were used throughout library preparation.

### Single Molecule Footprinting of crosslinked chromatin

ES cells grown under standard conditions were washed once with PBS and supplemented with fresh ES medium 2h prior to fixation. After 2h, cells were washed twice with room temperature PBS, all liquid was carefully removed and 10ml of crosslinking medium (1% formaldehyde in DMEM) was added to plates, followed by a 10 minutes incubation on a shaker at room temperature. A replicate plate was spared from fixation and trypsinized to count the number of cells per plate. After 10 minutes, crosslinking was stopped by addition of 500µl of 2.5M Glycin per plate, followed by a 10 minutes incubation on a shaker at 4 degrees. All liquid was removed and 1ml ice cold PBS was added, followed by collection of cells using scraping. Fixed material was centrifuged at 600g for 5 mins at 4 degrees, resuspended in 1ml detergent containing Paro buffer (0.25% Triton X-100, 10mM Tris pH 8, 10mM EDTA pH 8, 0.5mM EGTA) and incubated on ice for 10 minutes. Material was spun down again, resuspended in SMF wash buffer (see above), followed by another centrifugation resuspension in 1ml SMF wash buffer. Using cell count measures, material equating to 0.5 million cells were aliquoted into a new tube, spun down and resuspended in 300µl 1x GC Buffer (50mM Tris (pH 8.5), 50mM NaCl, 10mM DTT) and transferred to a Bioruptor Pico tube, followed by sonication at Bioruptor Pico instrument for 10 cycles at 15 secs ON / 90 secs OFF settings. M.CviPI reaction mixture (50mM Tris (pH 8.5), 50mM NaCl, 10mM DTT, 640µM SAM, 0.3M Sucrose) was prepared and used to equilibrate a Slide-A-Lyzer MINI Dialysis Device (0.5 mL, cutoff 3.5K MW, Thermo 88400). Sonicated chromatin was loaded into the dialysis cuvette, which was set up in a 15ml Falcon tube filled with M.CviPI reaction mixture. 100U of M.CviPI was added to the dialysis cuvette, followed by incubation at a 37 C water bath for 4h. Reaction was replenished with 20U of M.CviPI and 4µl of 32mM SAM after every hour. After 4h (3 replenishments), 10mM MgCl2 was added to the reaction mixture, SAM was replenished and 40U of S.ssI was added to the dialysis cuvette, followed by 2 hours of incubation at 37 degrees, where SAM and S.ssI were replenished after 1h. After 6 hours of total footprinting, reactions were stopped by addition of a SDS-containing buffer (20mM Tris, 600mM NaCl, 1%SDS 10mM EDTA), RNase A and incubated at 60 degrees over-night. Proteinase K was added to the samples, which were then incubated at 65 degrees for 2 hours, followed by phenol/chloroform extraction of the foot-printed DNA.

### Amplicon bisulfite sequencing

A custom R script was used to design primers against in silico bisulfite converted templates, using Primer3 with slight modifications. Products ranged from 200 to 500bp in size with majority of amplicons being over 450bp. Primers were commercially synthesized and reconstituted in 96-well plates. DNA material obtained from one footprinting reaction (1µg) was bisulfite converted following standard Epitect bisulfite conversion kit protocol (Qiagen 59124). Entire material of one conversion was equally distributed to a 96-well plate and amplified via PCR using KAPA HiFi Uracil+ Kit (Roche) in a total volume of 16µl, with 625nM primers (forward and reverse combined) under following cycling conditions: 95C for 3min, 35cycles of 20sec at 98C, 30sec at 56C, 60sec at 72C, followed by 5mins at 72C. 10µl of each reaction was pooled together and subjected to 0.8x AMPureXP bead purification (Beckman Coulter A63880). 1µg of purified DNA was used for each amplicon bisulfite sample and sequencing libraries were prepared using the NEBNext Ultra II Kit (E7645). Up to 12 libraries were multiplexed using NEBNext Multiplex Oligos (E7335) and sequenced on a MiSeq instrument, generating 250bp paired-end reads. All targeted amplification experiments have been performed in duplicates.

### siRNA transfection and TF knockdown

Dharmacon ONTARGETplus siRNAs against CTCF, NRF1, MAX and NFYA were ordered as SMARTpools, consisting of 4 different siRNA sequences targeting each mRNA, and resuspended in sterile water. As control, ON-TARGETplus non-targeting control pool (Horizon Discovery, D-001810-10) was used at the same concentration. Following guidelines of the manufacturer for ES cells, DharmaFECT 1 (Horizon Discovery, T2001) was used as transfection reagent. Transfection was performed in gelatinized 6-well plates (Falcon #353934) seeded with 250.000 DNMT TKO ES cells, according to the instructions of the manufacturer, with a final siRNA concentration of 25nM and a final DharmaFECT volume of 4ul in each well with a total volume of 2ml Opti-MEM reduced serum medium (ThermoFisher 51985034). 24h post-transfection, siRNA medium was exchanged with standard ES medium. After a recovery period of 24h in standard medium, cells were supplemented again with the transfection mixture for further knockdown of the targets. After a total treatment of 72h, cells were pooled in two independent technical replicates, counted and aliquoted for footprinting according to standard SMF procedure as well as RNA- and protein isolation.

### Confirmation of siRNA knockdowns via qPCR and Western blots

siRNA transfected cells were pelleted and resuspended in TRIzol reagent (ThermoFisher 15596026) or Protein Extraction Buffer (1% Triton-X-100, 150mM NaCl, 50mM Tris pH=8) for RNA and protein purification, respectively. Upon purification, RNA concentration was measured used Qubit RNA High Sensitivity Kit and 500ng RNA was reverse transcribed using SuperScript IV (ThermoFisher 18091050) with random hexamer priming. cDNA was diluted 20-fold in PCR-grade water and subjected to real-time quantitative PCR (qPCR) using SYBR Green PCR Master Mix (ThermoFisher 4309155) with gene specific, intron-spanning primers. Primers were check for quantitative range with dilution series, and knockdown data was analyzed using DeltaDeltaCT method. Western blots were used to confirm the knockdowns on protein expression level. 5ug total protein from transfected cells were subjected SDS-PAGE (12% Gels) and transferred to a nitrocellulose membrane at 150V for 1h. Transfer was confirmed with Ponceau staining and blots were incubated with primary antibodies overnight. NRF1 antibody (Abcam ab55744) was used at 1:1000 dilution and CTCF antibody (Sigma 07-729) was used at 1:2000 dilution in TBS-T. Blots were washed twice, incubated with secondary antibodies for 1h at room temperature and imaged using at a Bio-Rad gel documentation system. Consequently, blots were washed with TBS-T, incubated with tubulin primary antibody for loading control (1:1000 dilution) and imaged in the same manner.

### Quantification and statistical analysis

#### Data alignment and methylation call

SMF data were processed as previously described (Krebs et al., 2017). Briefly, raw sequence files were pre-processed using Trimmomatic to remove Illumina adaptor sequences, remove low quality reads and trim low quality bases. The trimmed reads were then aligned using QuasR (using Bowtie as an aligner) (Gaidatzis et al., 2014) against a bisulfite index of the Mus Musculus genome (BSgenome.Mmusculus.UCSC.mm10). For other datasets (ChIP-seq, RNA-seq, MNase-seq), reads were aligned using QuasR against the Mus Musculus genome (BSgenome.Mmusculus.UCSC.mm10). Context independent cytosine methylation call was performed using QuasR. Custom R functions were developed to determine context dependent (CG, GC) average methylation. Methylation has been called genome wide for Cs covered at least 10 times.

#### Single molecule methylation call

Single molecule C methylation extraction was performed using QuasR (Gaidatzis et al., 2014). Custom R functions have been developed to determine nucleotide context and sort the molecules according to their methylation pattern using a molecular classifier.

#### Mapping of TF motifs

Position weight matrices (PWM) for vertebrates TFs were downloaded from the Jaspar database (Mathelier et al., 2015) and mapped over non repetitive regions of the mouse genome (BSgenome.Mmusculus.UCSC.mm10) using the matchPWM function of the Biostrings package. PWM with a score of 10 or above were retained for single molecule TF classification.

### Data analysis

All analyses were performed using R-Bioconductor. Ad hoc R scripts are available upon request. Comparison with external datasets Data collection was performed using the qCount function of QuasR (Gaidatzis et al., 2014). Reads were counted in a window around each TF motif [-100:100] using the qCount function of QuasR (Gaidatzis et al., 2014) and enrichment over input were derived. Correlations were calculated on log2 transformed data.

#### Identification of dependencies between TFs

For each analyzed TF pair, states frequency was computed (Bound-Bound, Bound-Unbound, Unbound-Bound, Unbound-Unbound) and to build a contingency table. A Fischer’s exact test was performed on the contingency table representing each TF pair, with the null hypothesis that the binding states of two TFs are not dependent (odds ratio equals 1). The output of this test is an odds ratio, representing the likelihood that TF1 and TF2 will be in the same state (e.g. both bound) over the likelihood that TF1 and TF2 will be in different states. Fischer’s test also calculates an associated p-value with the odd’s ratios. Here, we report the odd’s ratios of TF pairs as a proxy for dependency in their binding. The co-bound fraction was calculated for each TF in the pair individually by dividing the number of co-bound molecules by the total number of bound molecules by the TF.

## Acknowledgements

The authors are grateful to Feris Jung, Leslie Horner, Christiane Wirbelauer for technical assistance, Bernd Klaus and Lukas Burger for technical support for statistics. The authors would like to thank Charles Girardot and the Genome Biology Computational Support. The authors would like to thank Ralph Grand, Duncan Odom and members of the Krebs laboratory for helpful discussions and/or comments on the manuscript. Authors are thankful to Veronique Juvin (SciArtWork) for support with artwork and Guy Riddihough (LifeScienceEditors) for support in manuscript editing. Research in the laboratory of A.R.K is supported by core funding of the European Molecular Biology Laboratory, Deutche Forschung Gemenischaft (KR 5247/1-1) and a Swiss National Fund Ambizione grant (PZOOP3_161493). Salary of D.I was supported by a Swiss National Fund Ambizione grant (PZOOP3_161493). Research in the laboratory of D.S. is supported by the Novartis Research Foundation, the European Research Council (ERC) under the European Union’s Horizon research and innovation program (Grant agreement no. 667951) and the Swiss National Sciences Foundation. The authors would like to thank Henriques Laboratory for sharing the Overleaf bioRxiv template.

## Contributions

A.R.K designed the study. A.R.K and CS wrote the manuscript. C.S designed and performed the experiments with the help of R.K, D.I, L.V and V.B for Illumina sequencing developments and library preparation. A.R.K and D.S supervised conduction of the experiments. C.S. and A.R.K analyzed the data. All authors discussed the results and commented on the manuscript.

