## Supplementary Figures for "Single molecule occupancy patterns of transcription factors reveal determinants of cooperative binding *in vivo*"

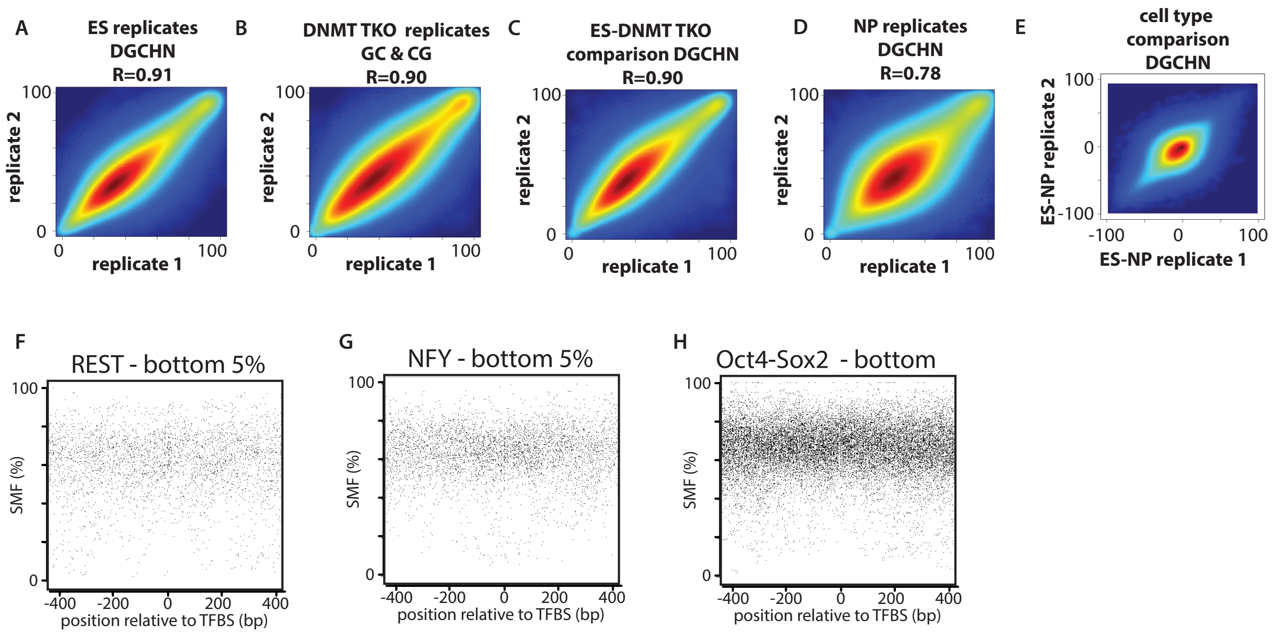


**Supplementary Figure 1: (A**) SMF replicates are highly correlated. Smoothed scatter plot representing the average genome wide methylation in GpC context (DGCHN) for the two biological replicates in mouse embryonic stem cells (mESC). **(B)** dSMF replicates are highly correlated. Smoothed scatter plot representing the average genome wide methylation in GpC or CpG context for the two biological replicates in DNMT triple knock out (TKO) mouse embryonic stem cells. **(C)** Accessibility profiles of WT and TKO mESC are highly correlated. Smoothed scatter plot representing the average genome wide methylation in GpC context (DGCHN) in WT and TKO mESC. (**D**) SMF replicates are highly correlated. Smoothed scatter plot representing the average genome wide methylation in GpC context (DGCHN) for the two biological replicates in mouse neuronal progenitors. (**E**) Methylation footprinting detects variation of accessibility between cell types. Smoothed scatter plot representing the average GpC methylation (DGCHN) difference between mESC and (NPs) compared between biological replicates. **(F-H)** Unbound regions lack discrete TF footprints in SMF data. Shown are the composite profiles of SMF signal at various (F, G, H) unbound TF motifs (bottom 10% of the respective TF ChIP-seq). Shown is the footprinting frequency (1-methylation [%]) of individual cytosines (black dots).

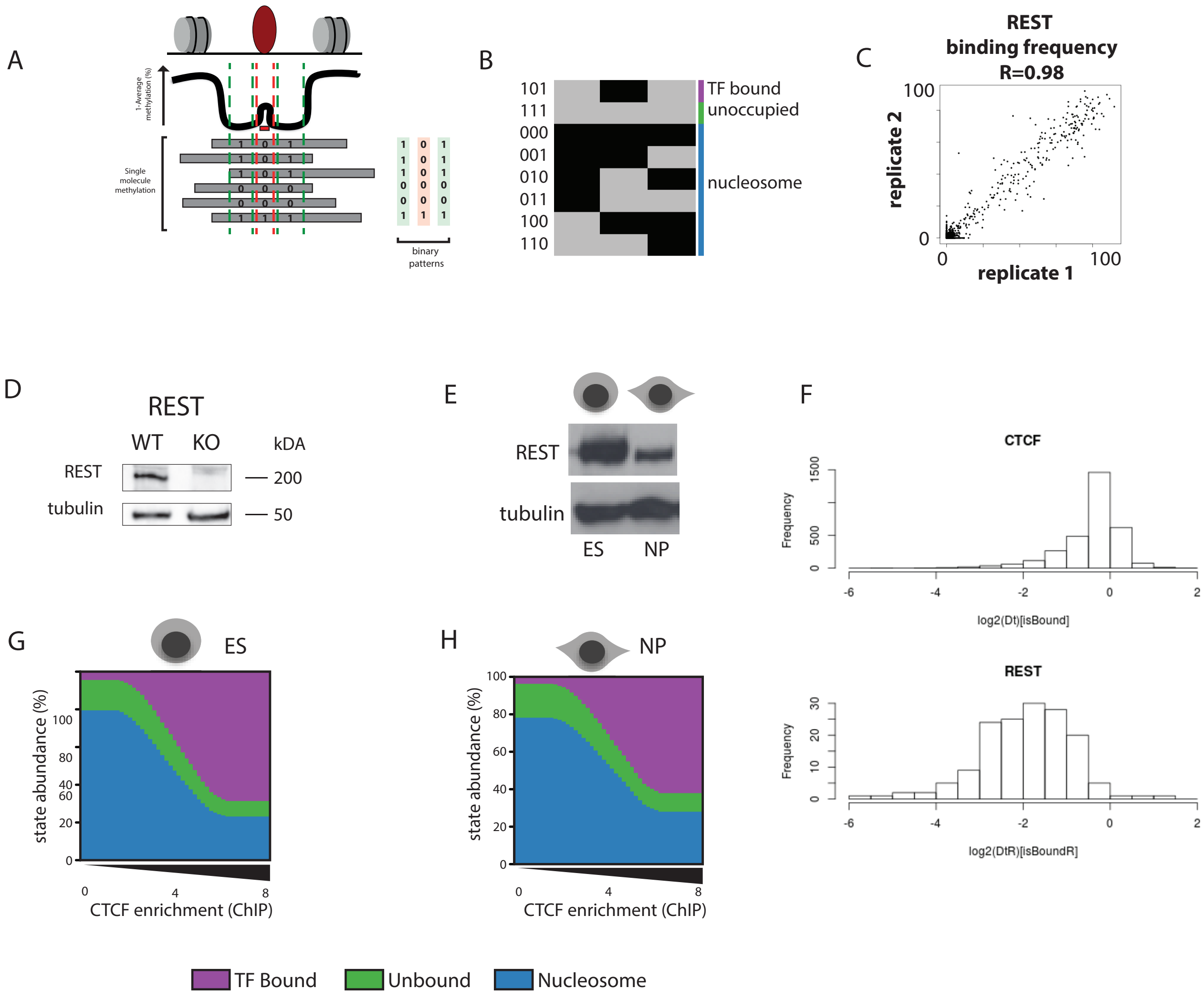


**Supplementary Figure 2: (A)** Schematic representation of the relative position of the three bins used for collecting TF footprints at the single molecule level. Color bars representing the position of each collection bin were overlaid on a typical average TF footprint. Bins were positioned to capture unoccupied molecules (green), the footprints created by a TF (purple) or a nucleosome (blue). **(B)** Schematic representation of the footprint patterns used to define each state (methylated Cs – accessible – light grey, unmethylated Cs - protected - black). States were named according to the type of occupancy expected to at these positions. **(C)** Quantification of REST binding frequencies in mESC is reproducible across biological replicates. Scatter plot representing the binding frequencies at all REST motifs covered. **(D)** Validation of the depletion of REST in knock out mESC. Western blot comparing REST protein levels in WT and REST KO mESC. **(E)** REST expression is reduced during neuronal differentiation. Western blot comparing REST protein levels in mESC and in vitro derived NPs. **(F)** Reduction of the REST binding frequencies in neuronal progenitors. Histogram depicting the changes in CTCF (upper panel) and REST (lower panel) binding frequencies during differentiation of mESC to NPs. **(G-H)** Distribution of state frequencies in mESC (G) and neuronal progenitors (NP) (H) as a function of CTCF occupancy as determined by ChIP-seq. Cumulative bar plot depicting the distribution of state frequencies. TFBS were binned based on CTCF enrichment in mESCs (log2 ChIP-seq), and the median frequency of each state was calculated within each bin. The frequency of each state is color coded according to the legend below.

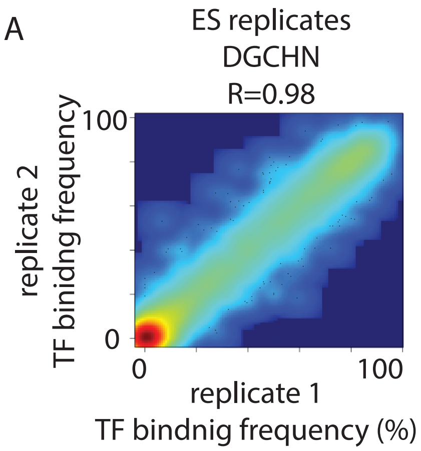


**Supplementary Figure 3: (A)** Quantification of TF binding frequencies in mESC is reproducible across biological replicates. Smoothed scatter plot representing the binding frequencies at all TF motifs covered.

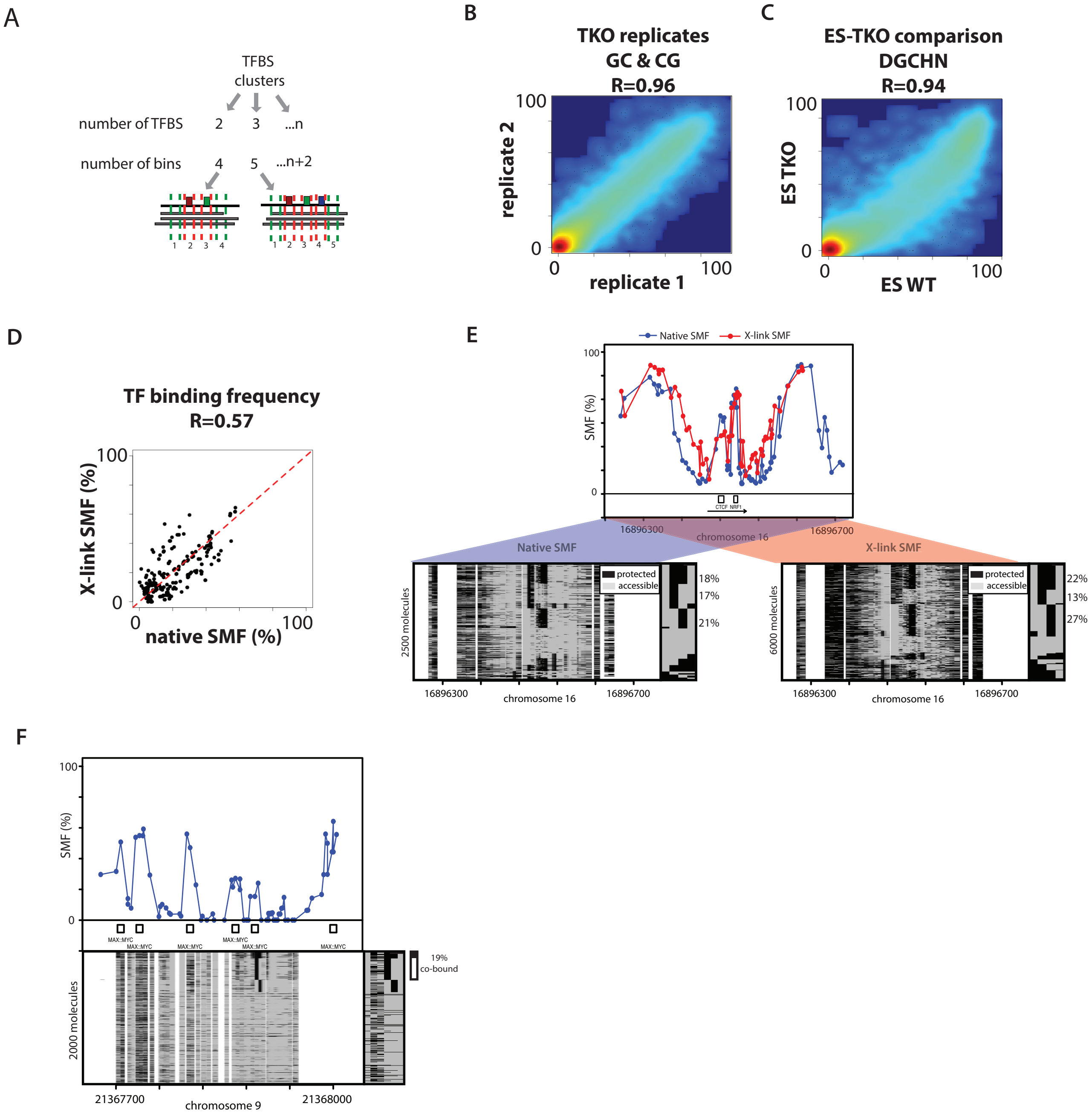


**Supplementary Figure 4: (A)** Schematic representation of the strategy used to classify reads at TF clusters. Loci are classified based on the number of TF motifs (n) present within the TF cluster. n+2 bins are positioned to capture the footprint around each motif (red) and the accessibility status in the region flanking the cluster (green). **(B)** Quantification of TF binding frequencies is highly reproducible across biological replicates. Smoothed scatter plot representing TF binding frequencies in two biological replicates of TKO mESC treated with CG and GC methyltransferases. **(C)** TF binding frequencies are highly correlated between WT and TKO mESC, suggesting that absence of DNA methylation has only marginal effects on the studied TF binding events. Smoothed scatter plot representing TF binding frequencies in WT and TKO mESC. **(D)** TF binding frequencies comparison between native SMF and X-link SMF. Scatter plot depicting the TF binding frequencies of TFs when performing footprinting on chromatin extracted after 10 min formaldehyde crosslink compared to standard SMF performed on permeabilized nuclei. The similarity between the methods confirm that the binding detected by native SMF reflect the binding behavior *in vivo*. **(E)** Single-locus example of the footprints detected using native SMF (top panel, blue dots connected by a blue line) or X-link SMF (red dots connected by a red line). Same representation as in Figure 4E. Very similar frequencies of co-occupancy are measured with both methods. **(F)** Single-locus example of a region bound by a Max-Myc dimer. Analysis of the degree of protein co-occupancy within the Max-Myc dimeric motif. Shown are average methylation levels (top panel, blue dots connected by a black line) and single-molecule stacks measured by targeted amplicon bisulfite sequencing (bottom left), sorted into four states using the co-occupancy classification algorithm (methylated Cs, accessible, light gray; unmethylated Cs, protected, black). The states heatmap (bottom right, binarized heatmap) depicts the occupancy states for each of the protein analyzed in the pairs. The percentages of molecules where individual or co-binding are observed is indicated on the right side of the plot. The fraction of co-bound molecules within all bound molecules is indicated in the sidebar.

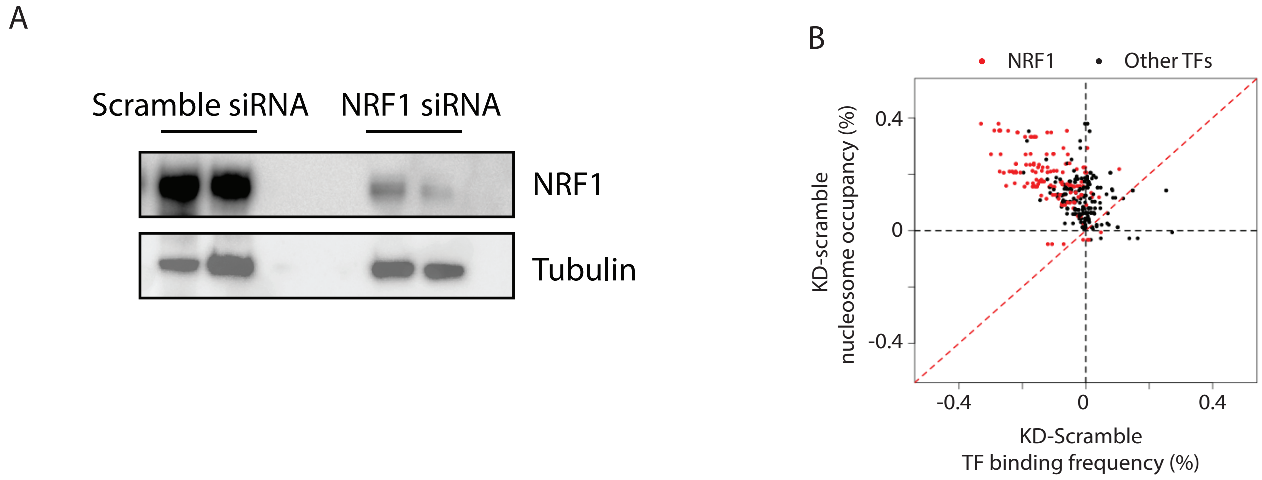


**Supplementary Figure 5: (A)** NRF1 knock down (KD) leads to reproducible reduction of protein concentration. Western blot comparing NRF1 protein levels in mESC treated with a Scramble and a NRF1 siRNA. **(B)** The decrease in TF binding frequency is tightly coupled with an increase in nucleosome occupancy at the NRF1 bound regions. Scatterplot depicting the difference in TF binding against the difference in nucleosome occupancy up in mESC treated with a Scramble or a NRF1 siRNA. Red dots correspond to the occupancy at NRF1 motifs while black dots represent any other motif.
